# Probabilistic Brain MR Image Transformation Using Generative Models

**DOI:** 10.1101/2024.11.16.623969

**Authors:** Sepideh Rezvani, Saeed Moazami, Christina J. Azevedo, Assad A. Oberai

## Abstract

Brain MR image transformation, which is the process of transforming one type of MR image into another, is a critical neuroimaging task that is needed when the target image type is missing or corrupted. Accordingly, several methods have been developed to tackle this problem, with a recent focus on deep learning-based models. In this paper, we investigate the performance of the conditional version of three such probabilistic generative models, including conditional Generative Adversarial Networks (cGAN), Noise Conditioned Score Networks (NCSN), and De-noising Diffusion Probabilistic Models (DDPM). We also compare their performance against a more traditional deterministic U-Net based model. We train and test these models using MR images from publicly available datasets IXI and OASIS. For images from the IXI dataset, we conduct experiments on combinations of transformations between T1-weighted (T1), T2-weighted (T2), and proton density (PD) images, whereas for the OASIS dataset, we consider combinations of T1, T2, and Fluid Attenuated Inversion Recovery (FLAIR) images. In evaluating these models, we measure the similarity between the transformed image and the target image using metrics like PSNR and SSIM. In addition, for the three probabilistic generative models, we evaluate the utility of generating an ensemble of predictions by computing a metric that measures the variance in their predictions and demonstrate that it can be used to identify out-of-distribution (OOD) input images. We conclude that the NCSN model yields the most accurate transformations, while the DDPM model yields variance results that most clearly detect OOD inputs. We also note that while the results for the two diffusion models (NCSN and DDPM) are more accurate than those for the cGAN, the latter was significantly more efficient in generating multiple samples. Overall, our work demonstrates the utility of probabilistic conditional generative models for MR image transformation and highlights the role of generating an ensemble of outputs in identifying OOD input images.

## 1 Introduction

Brain MR imaging is considered the primary method for diagnosing and monitoring patients with many neurological diseases, as well as conducting research on the brains of healthy subjects. This mainly stems from the ability of MR images to provide high-quality details of intracranial tissues. Additionally, MR images are highly flexible in terms of image acquisition characteristics. In other words, by tailoring the MRI sequence, a variety of image types can be acquired. These images have a wide range of clinical and research applications because of their ability to provide different anatomical and pathological details.

In many instances, not all of the desired MR image types and tissue contrasts are available. There are several reasons for this. These include the physical condition of the subjects, the cost of additional acquisition time, corruption of data, or the time window for the required scans being closed. The non-availability of all MR image types can have significant consequences. It can mean the exclusion of a subject from a study because of just one missing image type. For studies that involve machine learning algorithms, this can lead to the loss of a large amount of useful training data. In a clinical setting, it can mean incorrect diagnosis due to missing data or unnecessary repetition of MRI scans. For all these reasons, it is useful to be able to generate one MR image type from another. This medical image synthesis process is often referred to as MR image transformation.

Generally, a wide range of methods have been developed in the literature for medical imaging applications. More recently, Deep Learning (DL)-based models have proven to be successful in performing various tasks in this field [1, 2]. This mainly stems from the inherent capability of DL-based models in capturing the complex and highly non-linear mapping required in image-to-image transformations. The reader is referred to review articles like [3, 4] that review studies that focus on transforming various medical images of one kind to another. This includes inter-modality (from one modality to another) and intra-modality (from one image type to another of the same modality) conversion. The transformation from MR to computed tomography (CT) images [5], MR to positron emission tomography images (PET) [6], PET to CT [7], and the reverse of these transformations [8] are examples of inter-modality conversion. On the other hand, the transformation from T2 MR to FLAIR MR image types and its reverse are examples of the intra-modality transformations. The focus of this work is on the intra-modality transformation of MR images, and in particular, on the conversion of structural MR images from one image type to another.

Numerous variants of neural network architectures have been developed for DL-based models in the medical imaging field. While earlier works mostly rely on deep convolutional and fully connected neural networks (CNN and FCNN) [9, 10], U-Net-based architectures have received more attention for medical imaging tasks in recent years [11, 12] due to their high performance in medical imaging tasks. Accordingly, we adopt this architecture for all models implemented in this work.

Besides the model architecture, several methods and frameworks have been introduced in the literature to perform image transformation. Direct inference supervised methods [13] can be considered the most straightforward DL-based algorithms, in which the model receives the input and directly provides the output. During the training, a loss function, usually in the form of the difference between the prediction and the target or ground truth, is minimized. The aim of this process is to provide predictions that are as close as possible to the target. For example, the authors in [14] use an encoder-decoder using dilation CNN and least absolute error (*L*_1_) to generate brain CT images from T1 MR images. In another work [15], an MRI-to-CT conversion model is implemented via several embedding blocks. The loss function consists of the least squares error (*L*_2_) between the ground truth and the intermediate layer outputs in addition to the final prediction. In this manuscript, we have also implemented a model with U-Net architecture and used the supervised learning approach as a reference to compare the performance of other models. This is discussed in Section 2.2.

More recently, a sub-class of DL-based models called generative models have received significant interest in performing a wide range of applications. As we will discuss in Section 2.3, these models aim to learn the underlying distribution of the data during the training process. Thereafter, they generate samples from the learned distribution. Among the variants of generative models, two major classes of algorithms, generative adversarial networks (GANs) [16, 17, 18] and diffusion models [19, 20], have demonstrated promising performance in image-to-image transformation tasks. For example, the authors in [21] discuss a conditional generative adversarial network for MR image transformation. In another work [22], the authors discuss the effectiveness of cyclic consistency loss in addition to adversarial loss, especially in the transformation of unregistered MR images. Moreover, the study presented in [23] investigates the performance of diffusion-based models in converting MR to CT images and compares it with other DL-based methods. The authors in [24] present SynDiff, which combines a conditional diffusion process with an adversarial projection scheme for faster image sampling during inference. In doing so, they also account for cyclic consistency and are therefore able to handle unpaired data. There are also works in the literature that focus on low-field MR and low-dose PET imaging [25]. For example, in [26], the authors investigate the feasibility of synthesizing high-quality 3T MR images using scans from low-field 64mT portable MR devices. They present the LoHiResGAN method with a compound loss function that incorporates structural similarity index and *L*_1_ terms in addition to adversarial loss term.

There are two broad perspectives of looking at the image transformation problem. The first perspective, which is the one that is commonly used, is deterministic. In particular, this approach involves solving a regression problem for a transformation that maps images of one type to another. In this case, the transformation itself is deterministic, and for one input, it generates a single instance of the output. We refer to this approach as the “deterministic” approach.

Another perspective, which is explored in this work, looks at samples from the two different types of images as samples drawn from a joint probability distribution. Thereafter, it trains a conditional generative model to learn this distribution and then draws samples from the conditional distribution conditioned on one image type. The key difference between this perspective and the deterministic perspective is that for a given input, this perspective allows for the possibility of multiple samples. Thus, the transformation of an input image is not deterministic; rather it is “probabilistic”. It leads to multiple samples that are all drawn from the same conditional distribution. A key advantage of this approach is that the ensemble of transformed samples allows for a quantitative assessment of the uncertainty in the transformed image which can then be used for downstream tasks such as assessing the reliability of a transformation.

There are instances in the literature that have focused on developing probabilistic transformations in other medical imaging tasks. For example, in [27], conditional generative models were used for the brain extraction task. It was shown that the uncertainty computed correlated with error in the brain extraction and could also be used to determine whether a given input image was out-of-distribution. In [28], the authors propose a generative segmentation model based on a combination of a U-Net with a conditional variational autoencoder to produce an ensemble of possible segmentations for lung abnormalities. They utilize a dataset with multiple target images per input image for the training process.

In this work, we discuss the application of three popular conditional generative models for transforming MR images. In particular, we utilize a modified version of the conditional Wasserstein generative adversarial network (cGAN) [29, 30, 31, 32]. We also consider a conditional version of the noise conditioned score network (NCSN) [33, 34, 35]. Finally, we also implement a conditional denoising diffusion probabilistic model (DDPM) model based on the work presented in [36], which is a conditional version of the algorithm discussed in [37]. We compare the performance of these probabilistic transformation algorithms with a reference deterministic algorithm.

In the following, in Section 2, we first discuss the two distinct formulations of the image transformation problem. The first is a regressing problem where we look for the transformation that minimizes the *L*_2_ error. The second is the problem of sampling from a conditional distribution, given that samples are from the joint distribution, which leads to a probabilistic transformation. In this section, we also detail the pre-and post-processing procedures, evaluation metrics, and the datasets used for training and evaluation. In Section 3, we extensively discuss the qualitative and quantitative results, compare the performance of the models, and demonstrate the advantages of the probabilistic transformation approach. Finally, in Section 4, we deliver the findings and remarks derived from our experiments in addition to some suggestions and directions for future work and potential extensions.

## 2 Materials and Methods

### 2.1 MR image transformation problem formulation

In this section, we investigate the MR image transformation problem from both deterministic and probabilistic perspectives. To do so, we assume that a dataset of *N* paired MR images from an input and an output image type is available. This dataset is denoted by 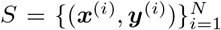, where 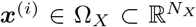 are images from the input image type, and 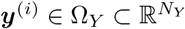 are images from the output image type. Here *N*_*X*_ and *N*_*Y*_ are the number of pixels in the output and input images, respectively. In the examples considered in this study, *N*_*X*_ = *N*_*Y*_ where each image is a two-dimensional axial image of the brain. It is further assumed that each image pair corresponds to the same physical slice and belongs to the same subject.

In the deterministic formulation of the problem, the aim is to solve a non-linear regression problem to estimate the pixel intensity of the output images from the inputs. To that end, as discussed in Section 2.2, we utilize a supervised model as a function approximator that captures the deterministic one-by-one mapping between the inputs and outputs.

We also define the MR image transformation task as a probabilistic inference problem and aim to solve it using the presented generative models. In practice, the dataset *S* is the same as in the deterministic approach. However, in a probabilistic framework, we treat the images from the input and output image types as random variables ***X*** and ***Y***, respectively. Then, the paired images of dataset *S* represent samples drawn from a joint density function *p*_***XY***_ (***x, y***). Using dataset *S*, we aim to train deep generative models that receive a new input image 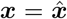 and can generate an ensemble of likely transformed generated images given the input image as 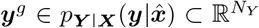. These generated samples can be used to compute any desired point statistic, including a) a mean image of the generated samples as a single best guess for the inferred transformed image and b) a standard deviation map of the generated samples that can be interpreted as the variation in the outputs or uncertainty of the models in their prediction.

As mentioned earlier, we aim to solve the deterministic formulation of the MR image transformation problem using a supervised learning model discussed in 2.2. We also solve the probabilistic version of this problem using three distinct types of generative models: conditional Wasserstein generative adversarial networks (cWGAN or cGAN), noise conditioned score network (NCSN), and denoising diffusion probabilistic models (DDPM) methods. These are described in detail in Sections 2.4, 2.5.1, and 2.5.2.

### 2.2 Deterministic supervised learning model

We implement a deterministic supervised regression model for MR image transformation using a direct inference framework. In this framework, we assume that there exists a function, 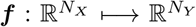, that directly transforms the pixel intensity of the input images to output images through a nonlinear transformation. Therefore, we model ***f*** using a neural network as a function approximator and denote it as 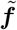. The training of this model involves minimizing a loss function that measures the difference between the output of the model 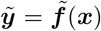 and the target or ground through ***y*** for all images of the training dataset (***x***^(*i*)^, ***y***^(*i*)^), *i* = 1 … *N*. This difference is formulated as the mean squared error (*L*_2_) as

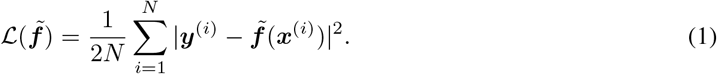

The optimized model is then given as 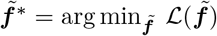. We refer to this model as the supervised or the direct model in this paper.

The motivation behind using this model is its convenience in implementation and training. As a result, we use this model as a reference to compare the performance of other models against it. We also note, in order to make a fair comparison, that U-Net neural network architecture is used for all presented models, i.e., the supervised model, the generator in cGAN, the score network in NCSN, and the denoiser in DDPM (see Section 4 for more implementation details.)

### 2.3 Generative models

As mentioned in Section 2.1, we aim to solve the probabilistic formulation of MR image transformation using a subclass of DL-based models called generative models. Generative models can generally be divided into two groups: unconditional (or pure) and conditional. Unconditional generative models are trained on a dataset of samples assumed to be drawn from an unknown probability density function, such as a large number of T1 MR images from different slices of different subjects. A fully trained unconditional model can generate new samples that also belong to the same underlying distribution, e.g., new T1 images in our example. One should note that new samples look like samples from training data, i.e., belong to the same distribution but are not exactly any of the data used during training.

In conditional generative models, the training is often done using paired data, such as a large number of pairs of T1 and T2 images where each pair belongs to the same subject and corresponds to the same slice location. These samples are assumed to be from the joint probability density function of these entities. Once trained, the model receives one input instance and generates samples from a probability density function conditioned on the provided input. Conditional generative models have significantly more applications because, unlike pure generative models, they provide the ability to have control over the generated outputs by determining the inputs (conditions).

It is noteworthy that some medical imaging problems, including MR image transformation, can also be solved using unpaired datasets. This is usually done using special considerations such as applying cyclic consistency loss terms [38]. However, if the paired data is available, conditional models often yield more accurate results [39]. Models that rely on unpaired data are necessary for applications where providing paired data is impossible or highly expensive. For example, in MR image harmonization [40], where collecting paired data involves traveling subjects, i.e., individuals who are required to visit multiple sites to provide scans from different devices.

Since conditional generative models generate samples from a conditional distribution, these samples can be used to compute important point statistics for the corresponding random variables. In our case, these include the pixel-wise mean and the pixel-wise standard deviation images for a given input image. This analysis is not possible when using a deterministic approach.

### 2.4 Conditional Wasserstein generative adversarial network (cGAN)

As shown in Fig. 1, the cGAN architecture consists of two deep neural networks, a generator ***g*** and a critic or discriminator *d*. The generator, ***g*** : Ω_*X*_ × Ω_*Z*_ ⟼ Ω_*Y*_, receives an image from the input image type as ***x*** and a random vector called latent variable 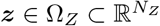. For any instance of ***z***, the generator can generate an output image as ***y***^*g*^ = ***g***(***x, z***). The distribution of the latent variable, *p*_***Z***_, is selected to be known and simple (a noise vector of size *N*_*Z*_ drawn from a standard multi-variable Gaussian distribution in our case). By passing a given input image ***x*** to ***g*** along with multiple instances of ***z*** drawn from *p*_***Z***_, the generator generates an ensemble of transformed images. The generated images can be considered samples of the likely transformed images drawn from a conditional distribution given the input image, 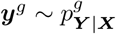.

**Figure 1:**
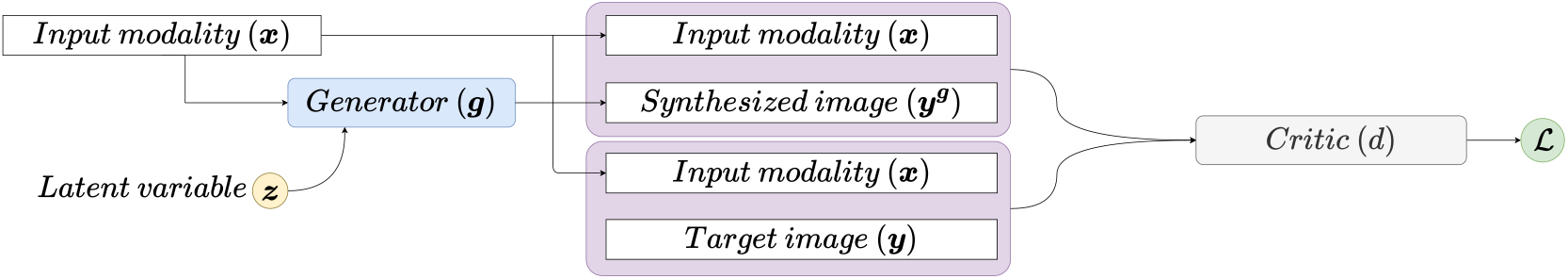
The conditional Wasserstein generative adversarial network (cGAN) architecture used in this work. The generator ***g*** receives the input image ***x***, and generates a synthesized image ***y***^***g***^ for any random latent vector ***z***. The critic *d* distinguishes between the generated synthesized image ***y***^***g***^ and target image ***y*** paired with ***x***. The training process involves updating ***g*** such that its outputs are indistinguishable from real images by *d* and updating *d* to become better in distinguishing the generated images from real ones.

The critic *d* : Ω_*X*_ *×* Ω_*Y*_ ⟼ ℝ receives a pair of images and is responsible for distinguishing between the pairs that are from the true dataset, i.e., (***x, y***) ∼ *p*_***XY***_, and those pairs where the output image type is generated by the generator network, i.e., (***x, y***^***g***^), where 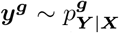. This process is done by training the critic to yield larger values for images from the dataset (true) and smaller values for the images generated by the generator (fake). This is achievable via the Wasserstein GAN adversarial loss function [30] as the expectation of the difference between the values of the critic for true and generated images

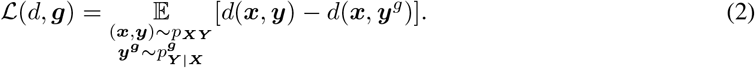

Then, the min-max optimization problem is solved concurrently to train the optimal generator and critic,

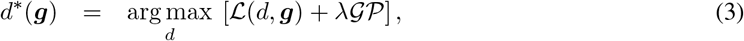

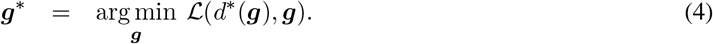

We note that the 𝒢𝒫 term in 3 is the gradient penalty [31], with *λ* hyper-parameter coefficient, and is defined as

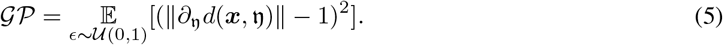

Intuitively, Equation 3 defines the optimal critic *d*^*^ as the function that produces the highest value for the loss function defined in Equation 2 for a given value of the generator. The gradient penalty term in 5 is used to enforce the 1-Lipshitz constraint on the critic by penalizing gradients with norms larger than one (see [32]). The gradients are calculated at a weighted value of the actual and generated images of the output image type given by 𝔶 = *ϵ****y*** + (1 −*ϵ*)***y***^***g***^ where *ϵ* ∼ 𝒰(0, 1). The loss function evaluated at the optimal value for the critic (that is the term ℒ (*d*^*^, ***g***)), provides a measure of Wasserstein-1 distance between the true joint distribution and its generated counterpart. Minimizing this distance by training the generator, which is achieved in Equation 4, leads to a ***g***^*^ that ensures that the true and generated joint distributions are as close as possible. This, in turn, implies that the generated conditional distribution 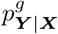 is close to the true conditional distribution *p* _*Y* |*X*_. Consequently, using any given image ***x*** as input to the generator, and then instances of the latent vector through the fully trained generator ***g***^*^ is equivalent to generating multiple samples from the true conditional distribution *p*_***Y*** |***X***_. It is noteworthy that this is a computationally efficient process as the latent variable is drawn from a known distribution with relatively small dimensionality compared to the images, *N*_***z***_ ≪ *N*_***X***_ = *N*_***Y***_, and each sample requires only one forward pass of the generator to be calculated.

### 2.5 Diffusion models

In this section, we provide a general intuition about diffusion models and then discuss the two major variants of these models in more detail in Sections 2.5.1 and 2.5.2. As illustrated in Figure 2, the general workflow of diffusion-based models consists of a forward noising process (from right to left) and a backward denoising process (from left to right). In the unconditional version of a diffusion model, a sample data point is drawn from the true initial distribution as ***y***_0_ ∼ *p*_0_(***y***), i.e., an image from the training dataset. Next, noise with varying scales (variance of 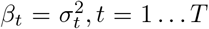) is added to ***y***_0_ for time steps *t* = 1 … *T*. This process is done until the image looks like pure noise, i.e., ***y***_*T*_ ∼ 𝒩 (***µ***, *β*_*T*_ 𝕀), where 𝕀 denotes the identity matrix. The noising process outputs form a sequence of noisy images ***y***_*t*_ for various *t* ∈ [1 … *T*]. The general idea is then to use these data to design a denoising process that can receive a noisy image ***y***_*t*_ ∼ *p*_*t*_(***y***) and generate a less noisy image ***y***_*t*−1_ ∼ *p*_*t*−1_(***y***) for any step *t*. This is done by training a deep neural network (the diffusion model) that receives an input image ***y***_*t*_ and a measure of *t* through timestep embedding [41]. The output of the model is an image that is used to generate ***y***_*t*−1_. The details of the processes can differ in variants of diffusion models and will be discussed subsequently for the two methods presented in this work. Eventually, this process is applied to an image drawn from the known simple distribution of pure noise ***y***_*T*_ ∼ 𝒩 (***µ***, *β*_*T*_ 𝕀) and after *T* denoising steps, *t* = *T* … 1, a sample from *p*_0_(***y***) is obtained.

**Figure 2:**
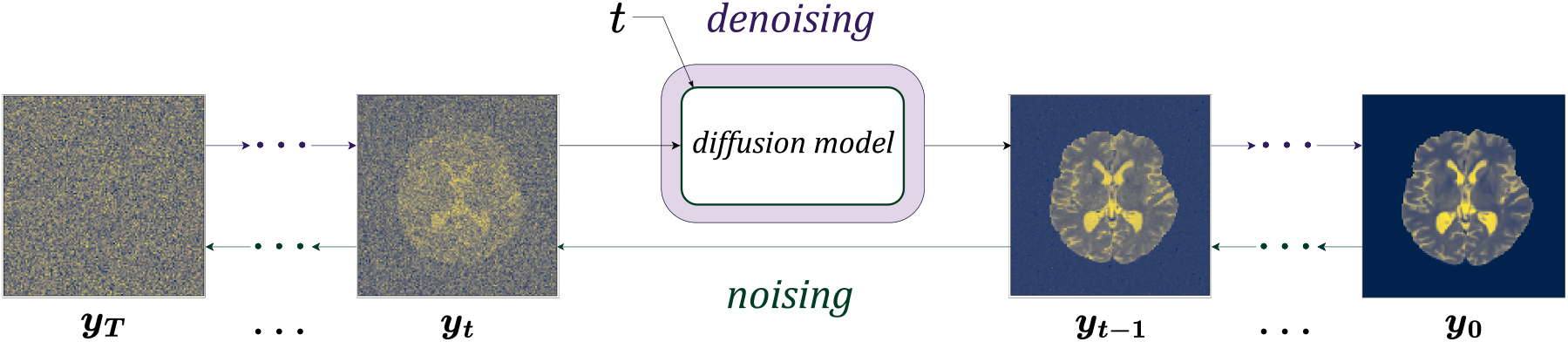
Forward noising and backward denoising processes in diffusion models. During training (from right to left) sample images go through several noising steps. This provides a series of images with different levels of noise. These images are used to train a model that receives a noisy image and time *t* and provides an output that is used to generate a less noisy image. After training, generating samples involves (from left to right) starting from a noise image and generating less noisy images, leading to the final generated image.

Intuitively, the process discussed above can be observed from two points of view. At the sample level, each image can be viewed as a particle in *N*_*Y*_ dimensional space. These are shown as dark points in Figure 3 for a simplified problem with *N*_*Y*_ = 2. Perturbing the samples by noise mimics the random walk or Brownian motion of individual particles in a diffusion process. At the population level, these particles collectively form samples drawn from the probability density function in *N*_*Y*_ dimensions. The contours of this distribution are plotted in Figure 3. The diffusion process represents the evolution of the data distribution *p*_0_(***y***) to a normal distribution 𝒩 (***µ***, *β*_*T*_ 𝕀) through *T* steps, as illustrated in Figure 3. This is equivalent to convolving the initial probability density function with a Gaussian kernel of increasing width. The process of generation or sampling moves in the opposite direction. That is, it evolves a sample from the Gaussian distribution *p*_*T*_ (***y***) to a sample from *p*_0_(***y***).

**Figure 3:**
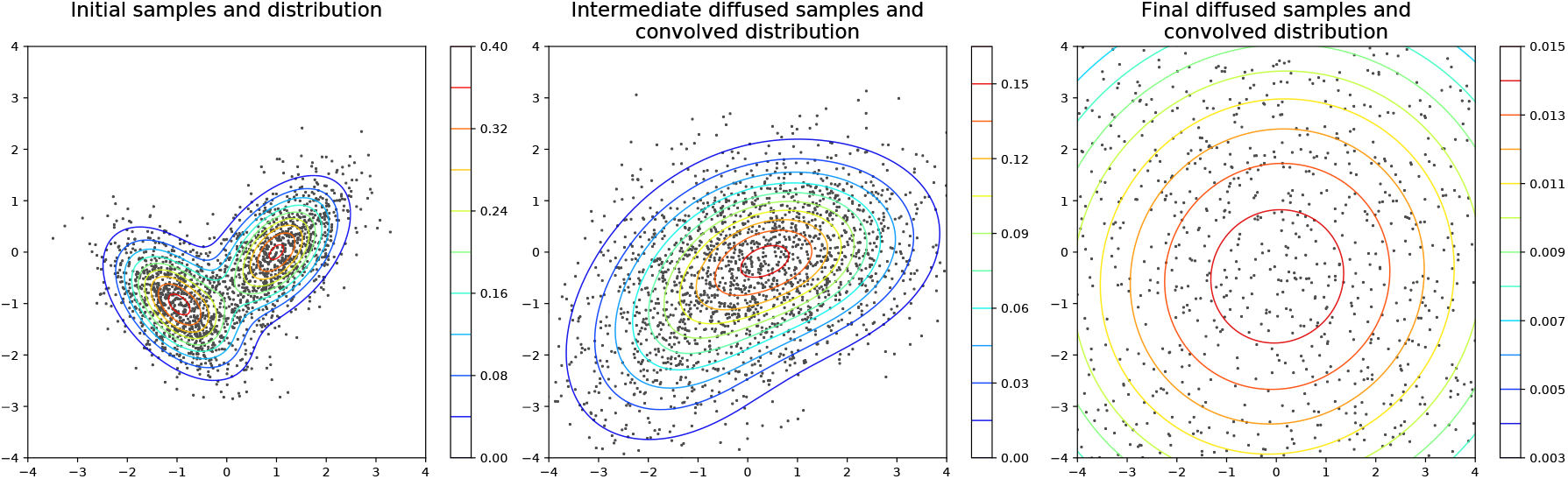
The diffusion process is demonstrated as probability density and samples. The figure on the left shows samples from an initial distribution. In the figure in the middle, these data points are diffused by adding a Gaussian noise. In the figure on the right, after extensive diffusing of the data points, the samples are now consistent with a Gaussian distribution with zero mean and a large variance.

In the conditional version, an extra image ***x*** is provided in addition to the input image for each step *t*, while ***y***_*t*_ undergoes the above-mentioned processes of noising and denoising. In this case ***x*** and ***y***_0_ = ***y*** are paired images from the dataset (***x, y***) ∈ *S*.

#### 2.5.1 Noise conditioned score networks (NCSN)

For the NCSN the noising step discussed in Section 2.5 involves adding a Gaussian noise with varying levels of variance *β*_*t*_ to the image drawn from the data distribution ***y***_0_ = ***y*** to generate intermediate images ***y***_*t*_, *t* = 1 … *T*, where,

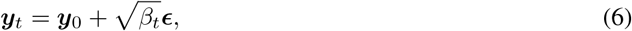

and 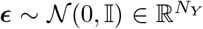 is a multi-variate Gaussian random vector. The series of *β*_*t*_ are hyper-parameters that are selected according to a schedule [34]. The training process in NCSN aims to directly learn a neural network model 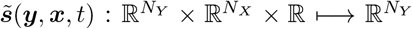 to approximate the score function (***s***(***y, x***, *t*) = log(*p*_***Y*** |***X***_ (***y x***, *t*)), *t* = 0 … *T* of the time dependent conditional distribution *p*_***Y*** |***X***_ (***y x***, *t*). This is achieved by defining an objective function

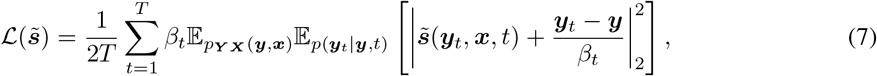

where the optimal score network model is defined as 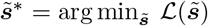.

The term within the square parenthesis in Equation 7 is similar in form to a regression loss function. Therefore, we recognize that the output of a fully trained score model provides an estimate of 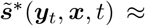 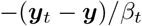. By rearranging this equation, we note that ***y*** 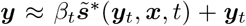. This implies that the estimated value of the score function, when multiplied by the variance *β*_*t*_, is what should be added to the perturbed image ***y***_*t*_ to generate the image without noise (***y***) at any step *t*. In practice, the denoising process in NCSN is performed iteratively through *T* steps of Langevin dynamics [34], where at any step, a less noisy image ***y***_*t*−1_ is generated using

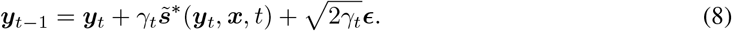

Here 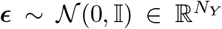 and *γ*_*t*_ = *λ*_*γ*_*β*_*t*_*/β*_1_ is the step size, *λ*_*γ*_ is a hyper-parameter, and *β*_1_ is the variance for the first time step (typically the smallest *β*_*t*_). In practice, the sampling process starts by setting ***y***_*T*_ ∽ 𝒩 (0, *β*_*T*_ 𝕀) and using this as input to the neural network for the score function along with input image ***x***. The initial image ***y***_*T*_ is denoised in *T* steps using Equation 8 to reach ***y***_0_, which is the transformed image generated conditioned on ***x***.

#### 2.5.2 Denoising diffusion probabilistic models (DDPM)

The noising process in the DDPM method also involves adding a tailored noise to the initial image ***y***_0_ as

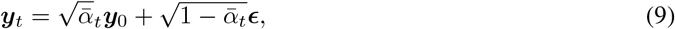

where *α*_*t*_ := 1 − *β*_*t*_ and 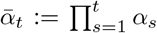 [42] and ***ϵ*** ∼ 𝒩 (0, 𝕀). In the DDPM method, unlike NCSN, the variances are limited to be between zero and one as *β*_*t*_ ∈ (0, 1), *t* = 1 … *T*, such that *β*_0_ *< β*_1_ *<* · · ·*< β*_*T*_. The noisy image ***y***_*t*_ generated through 9 and its counterpart from the initial distribution ***y***_0_ are used to train a denoiser function in the form of a neural network 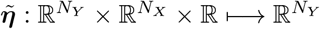. This network is trained using the loss function suggested by [42],

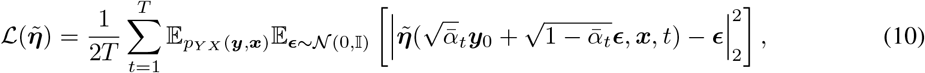

where the optimal denoiser function minimizes this loss function, 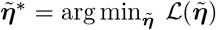. Similar to NCSN, the loss function in Equation 10 resembles a regression loss. Further, by comparing the term within the bracket of 10 and the noising process 9, we observe that 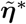 learns to provide an estimate of the noise that is added to the clear image to form the noisy image or 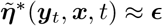.

The DDPM model used in this study is based on [37], which is an extension of an earlier work [42]. The training of this model involves minimization of a hybrid loss that contains an additional term so that the model also learns the sequence of noise scales 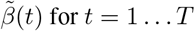, instead of accepting them as hyper-parameters set by the user (see Equations (15) and (16) of [37]). This network is denoted by 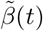 and it is trained concurrently with 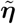.

The denoising process in the DDPM method is given by

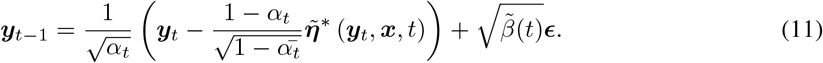

The sampling process begins by sampling ***y***_*T*_ ∽ 𝒩 (0, 𝕀), and using this and the image to transform, ***x***, as input to the denoiser network. The process in Equation 11 is then used to denoise and end with a new point sample ***y***_0_, which is the transformed image conditioned on the input image ***x***.

The NCSN and DDPM methods share many common features, including beginning with a sample drawn from a simple Gaussian distribution, iteratively denoising this sample using Langevin dynamics, and using a neural network that is trained using a simple regression loss. The key difference between the two methods is in the initial distribution from which samples are generated. In the NCSN method, this is a multivariate Gaussian distribution with zero mean and a very large variance, whereas in the DDPM method, it is also a multivariate Gaussian distribution with zero mean but with a variance of unity. The reader is referred to [43] for a unified description of these methods and their connections with the theory of stochastic differential equations.

### 2.6 Image transformation using generative models

We aim to investigate the ability of the generative models to provide multiple outputs for any given input. Accordingly, we discuss the procedure through which cGAN and two diffusion models (DDPM and NCSN) generate multiple output samples and how we use these in the inference process.

The inference process begins by loading a 2D axial image of the input brain image ***x*** as an input to the fully trained models. Thereafter, in the cGAN model, *n* samples of the latent vectors, ***z*** with dimension *N*_*Z*_ = 128 are used as input to the generator, which generates *n* images ***y***^(*i*)^, *i* = 1, …, *n* from the desired conditional distribution. For the diffusion model, the process is similar, however, the dimension of ***z*** is the same as that of ***y*** (*N*_***Z***_ = *N*_***Y***_ = 128^2^ in this work). These samples are treated as the initial state in the denoising process, that is ***y***_*T*_ = ***z***, whose termination yields the *n* generated images ***y***^(*i*)^, *i* = 1, …, *n*. We note that the sampling process in the diffusion models is rather expensive since it involves multiple steps. In our work, *T* = 1000 steps.

All three generative models generate an ensemble of images from the conditional distribution, which is used to compute a single pixel-wise mean image, 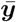,

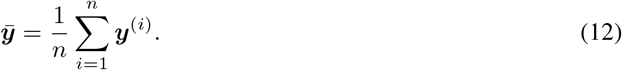

Similarly, an image of the pixel-wise standard deviation, ***y*′**, is calculated by

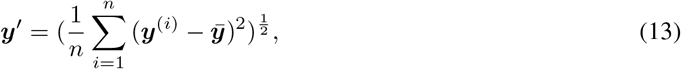

where the operations on the right hand side are interpreted as being pixel-wise.

### 2.7 Datasets

We use two public datasets with multiple image types, IXI [44] and OASIS [45], for training and evaluation of the proposed methods.

The IXI dataset consists of 600 MR images of three types: T1-weighted (T1), T2-weighted (T2), and proton density (PD). The scans are collected in three different hospitals in London from normal healthy subjects. Of these, 575 image sets passed QC (those with missing images and realignment issues were removed), from which 357, 74, and 144 images were used for the training, validation, and testing of the models. We balanced our datasets to have the appropriate number of scans from each hospital in the training, validation, and test datasets.

OASIS-3 is a public longitudinal dataset for normal aging and Alzheimer’s disease, consisting of T1, T2, and fluid-attenuated inversion recovery (Flair) images. Within this dataset, we used images from the participants that contain all image types scanned in the same MRI session. Out of 243 images, 156, 31, and 56 are used for training, validation, and testing of the model. In this paper, we use the same dataset splits for training and testing experiments of all models.

### 2.8 Pre-processing

For the IXI dataset, we used the Flirt module of Fsl [46] to realign the 3D T1 images to spaces of T2 images as they have the narrowest axial field of view. PD images do not require alignment to T2 images. As the user guide for Fsl suggests, this is done by first using T1 images as the reference as they have the highest resolution and then using the inverse transformation to provide T1 images in T2 space. For the OASIS dataset, we used T1 images as the reference and aligned the T2 and Flair images to them. We extracted the brains using T1 images as input using [27] and then applied the T1 masks to provide brain images for other images. Finally, we performed a min-max normalization.

### 2.9 Post-processing

We did not observe any systemic dissimilarity between the target and raw output of the Direct, cGAN, and NCSN models that could be further improved by applying an additional post-processing procedure. In contrast, DDPM models demonstrate isolated points with hyper or hypo-intense voxel intensities in their output images. This problem can be due to internal numerical issues as it also occurs as occasional undefined (value overflow) in the output, requiring special considerations. To handle this issue, we first identified these noisy voxels as outliers based on their higher pixel-wise standard deviation. Then, we used the pixel-wise median value of the generated samples instead of their mean to eliminate the effect of the outliers in these voxels. One should note that the generative capability of the DDPM model enables us to produce multiple samples to calculate the median required for eliminating noisy voxels.

### 2.10 Evaluation metrics

We used the Structural similarity index measure (SSIM) and peak signal-to-noise ratio (PSNR) to evaluate the overall quality and similarity of output images to the target images. The SSIM measures the similarity of two images, where every voxel from the target image is compared with the counterpart voxel in the prediction. In addition to being sensitive to pixel-wise differences, the SSIM metric can quantify the overall similarity of two images while accounting for their structure. This aspect is important for MR images where preserving the geometry of the organs through image transformation is essential. Therefore, we consider the SSIM the primary metric for the evaluation of the performance of the models. The PSNR metric quantifies the amount of signal and information between two images compared to their noise.

## 3 Results and Discussion

### 3.1 Image Transformation Results

In this section, we benchmark the performance of the models discussed in Sections 2.4, 2.5.2, 2.5.1, and 2.2 by applying them to a collection of images from the test datasets.

In Figures 4 and 5, we focus on the T1 to T2 transformation for axial slice images from the IXI and OASIS datasets. In these figures, in the first row, moving from left to right, we show the input T1 image, the target T2 image, and then the results from the Direct and the three probabilistic methods. In the second row, for each transformation method, we show the corresponding pixel-wise error, and in the third row, for each probabilistic method, we show the pixel-wise standard deviation.

**Figure 4:**
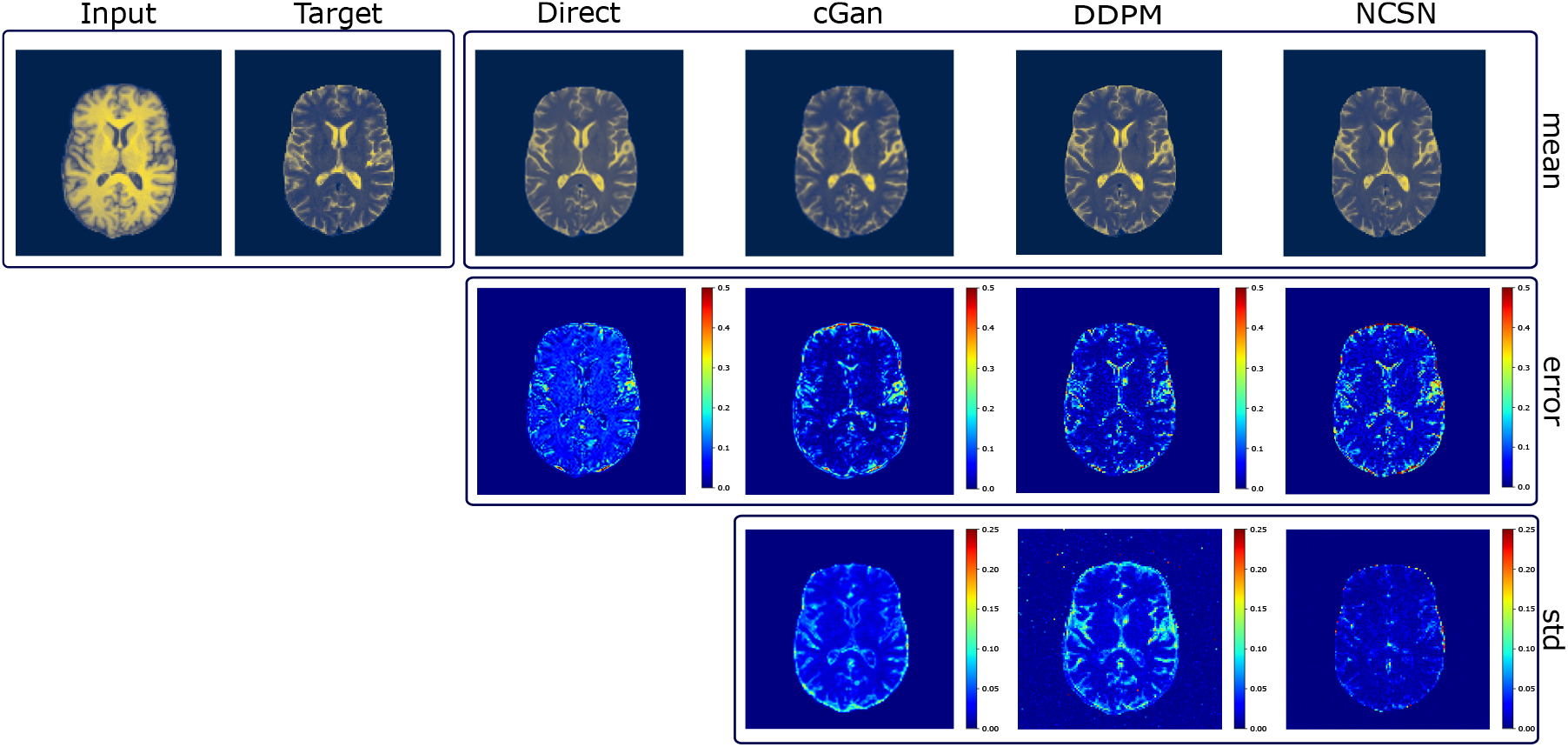
T1 to T2 MR image transformation for a sample slice of axial image from the IXI dataset. In the first row, the first two columns are the input image and ground truth (target). The remainder of the columns are output images from the Direct, cGAN, DDPM, and NCSN models. The second row is the pixel-wise absolute error for each model, and the third row is the pixel-wise standard deviation image for the generative models.

**Figure 5:**
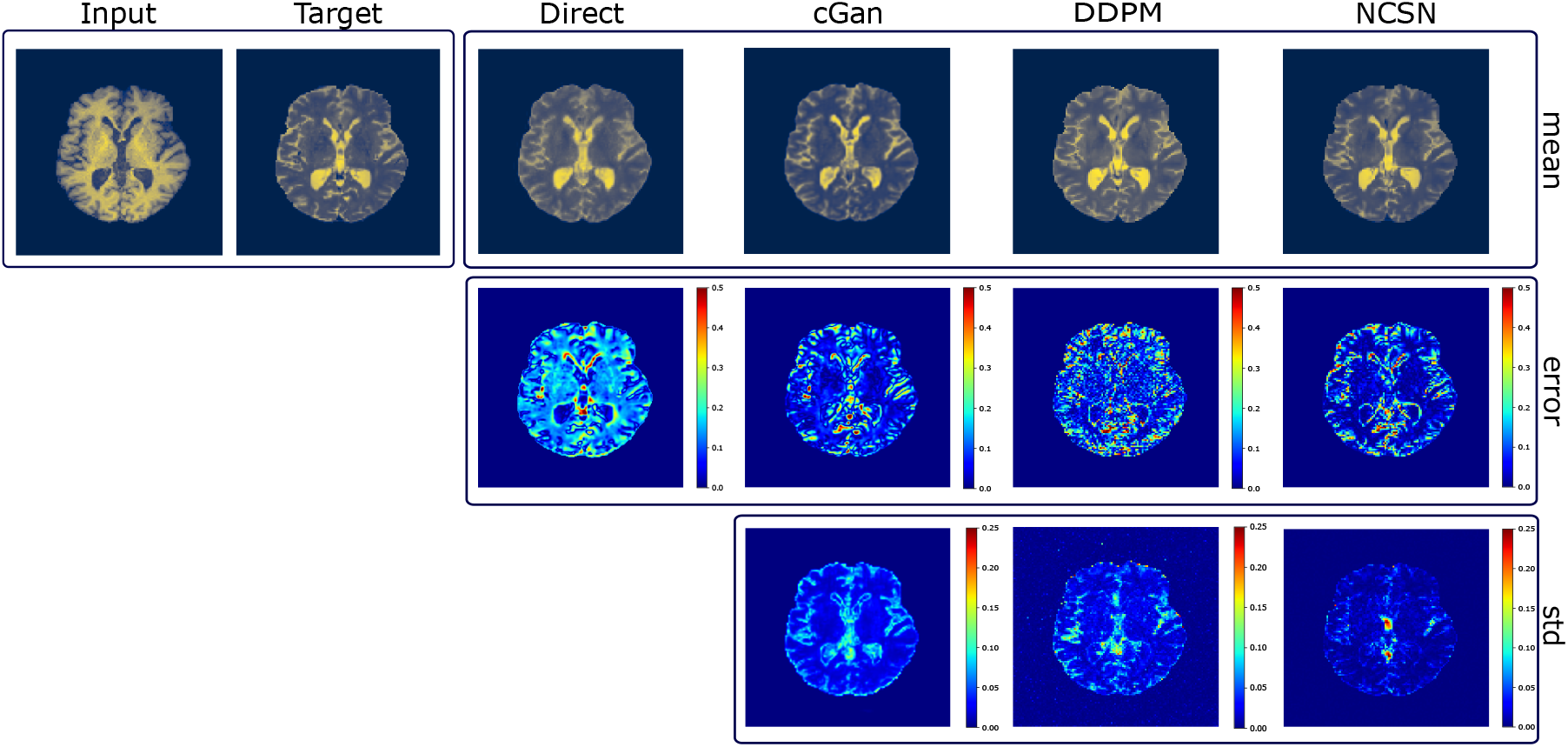
T1 to T2 MR image transformation for a sample slice of axial image from the OASIS dataset. In the first row, the first two columns are the input image and ground truth (target). The remainder of the columns are output images from the Direct, cGAN, DDPM, and NCSN models. The second row is the pixel-wise absolute error for each model, and the third row is the pixel-wise standard deviation image for the generative models.

In Figures 6 and 7, we demonstrate all possible transformations in a more compact format for the IXI and OASIS datasets, respectively. Each dataset contains three different image types, and we transform images from each of them to the other two. Consequently, in each figure, we present six results that are organized in six rows. From left to right, in the first and second columns, we present the input and ground truth or target images, respectively. We demonstrate the model outputs in the remaining columns. The third and fourth columns are the Direct model output and error, i.e., the absolute difference between the target and prediction. The fifth, sixth, and seventh columns contain the cGAN prediction, its pixel-wise error, and the predicted pixel-wise standard deviation. Columns 8-10 and 11-13 are the corresponding results for the NCSN and DDPM models, respectively.

**Figure 6:**
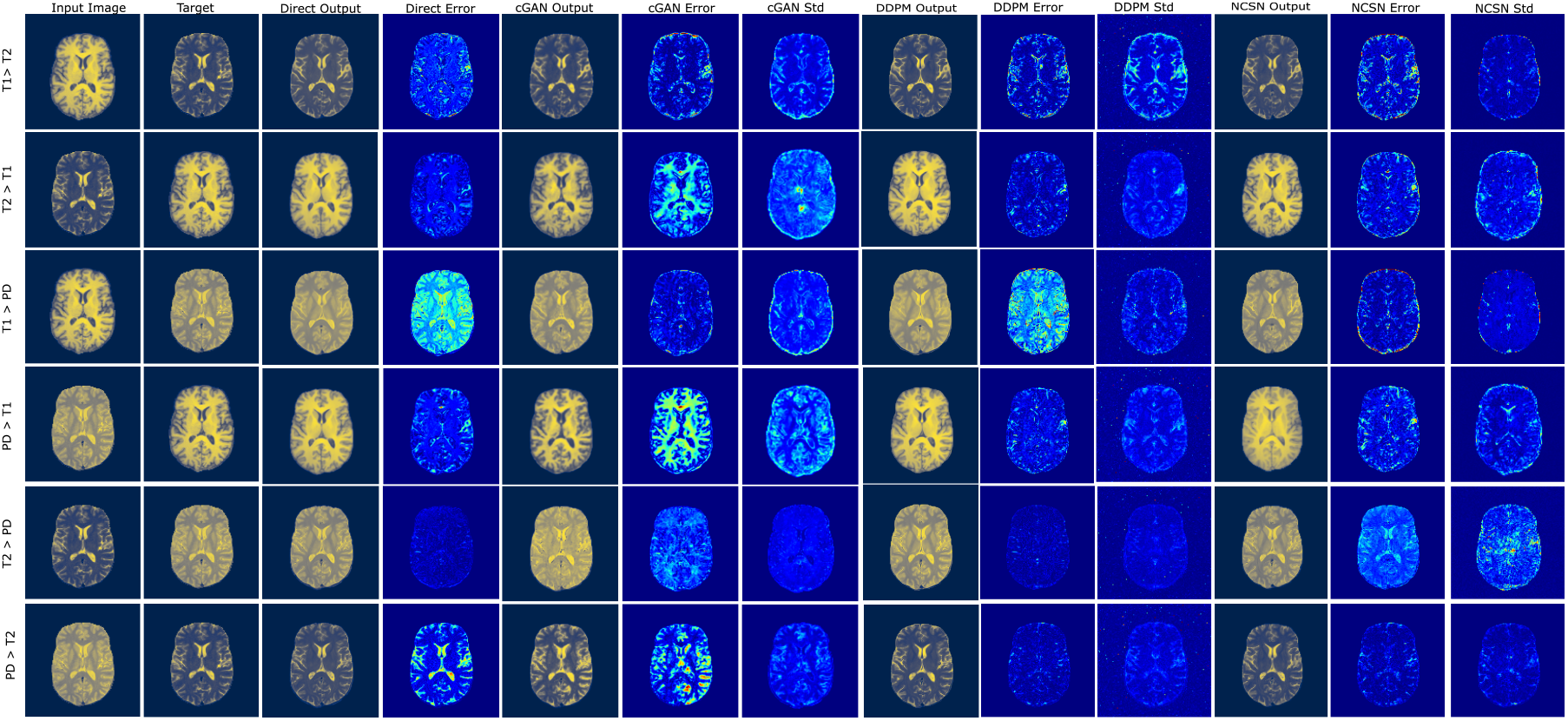
MR image transformation for an axial slice image from the IXI dataset. Column 1: input image; column 2: target image; columns 3-4: output and error for the direct model; columns: 5-7: output, error and standard deviation for cGAN; columns: 8-10: output, error and standard deviation for DDPM; columns: 8-10: output, error and standard deviation for NCSN.

**Figure 7:**
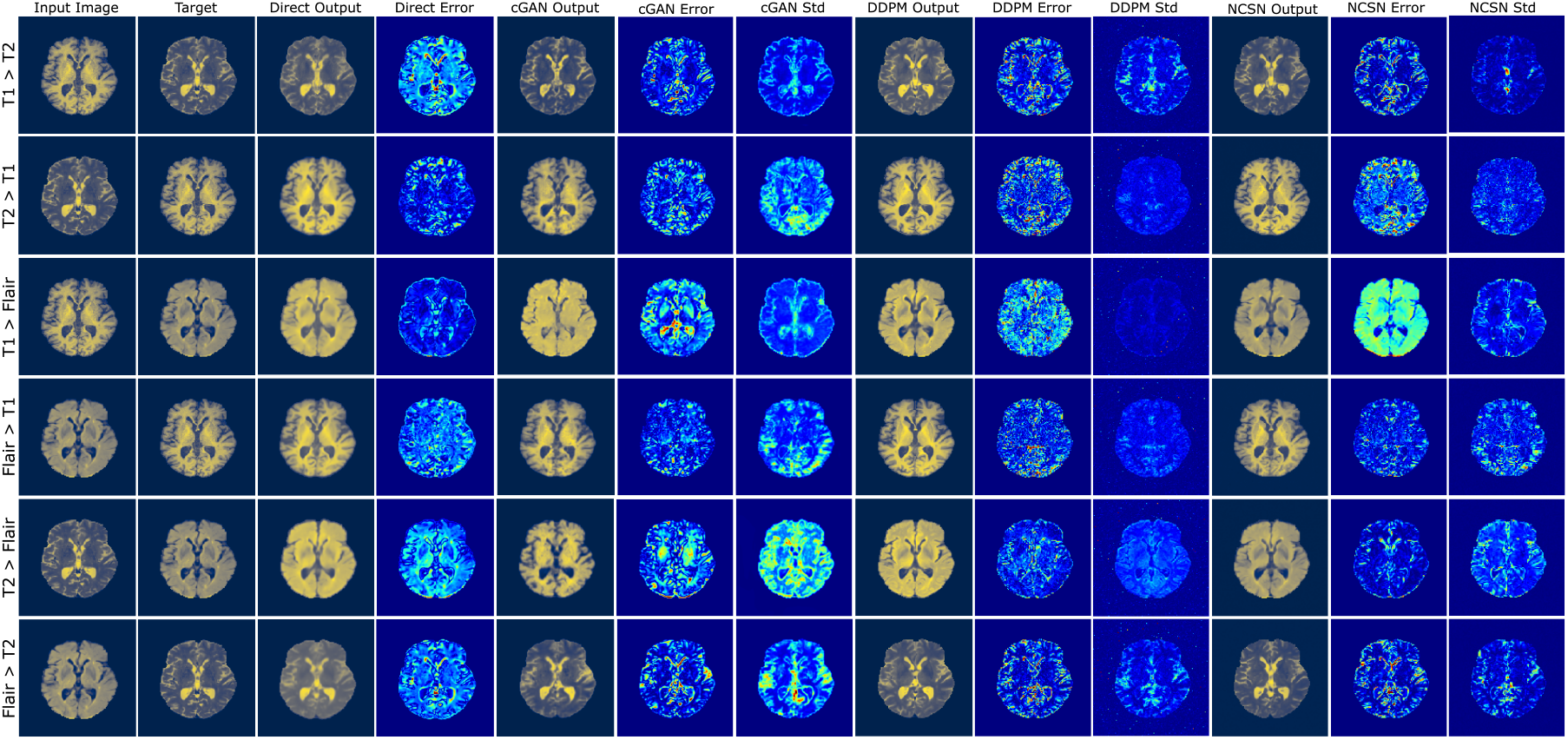
MR image transformation results for an axial slice image from the OASIS dataset. From left to right, the first two columns present the input image and ground truth (target). The third and fourth columns show the output and error for the Direct model, and the rest of the columns are the results (mean image, model error, and standard deviation image) for the remaining three models: cGAN, DDPM, and NCSN. As indicated, each row demonstrates the results for a different transformation.

We also illustrate the distribution of the calculated SSIM and PSNR metrics for the test subjects of the IXI and OASIS datasets in four plots in Figure 8. Each plot contains results for one metric (SSIM or PSNR) and one dataset (IXI or OASIS). Within plots, we consider six different transformations and demonstrate the performance of each transformation method in the form of a box plot. The rectangular extent of each box displays the interquartile range (IQR) where 50% of the values reside, and the whiskers depict the range of the values excluding the outliers. The outliers are defined as values more than 1.5 IQR below the 25^*th*^ percentile and 1.5 IQR above the 75^*th*^ percentile and are shown as markers in the plots.

**Figure 8:**
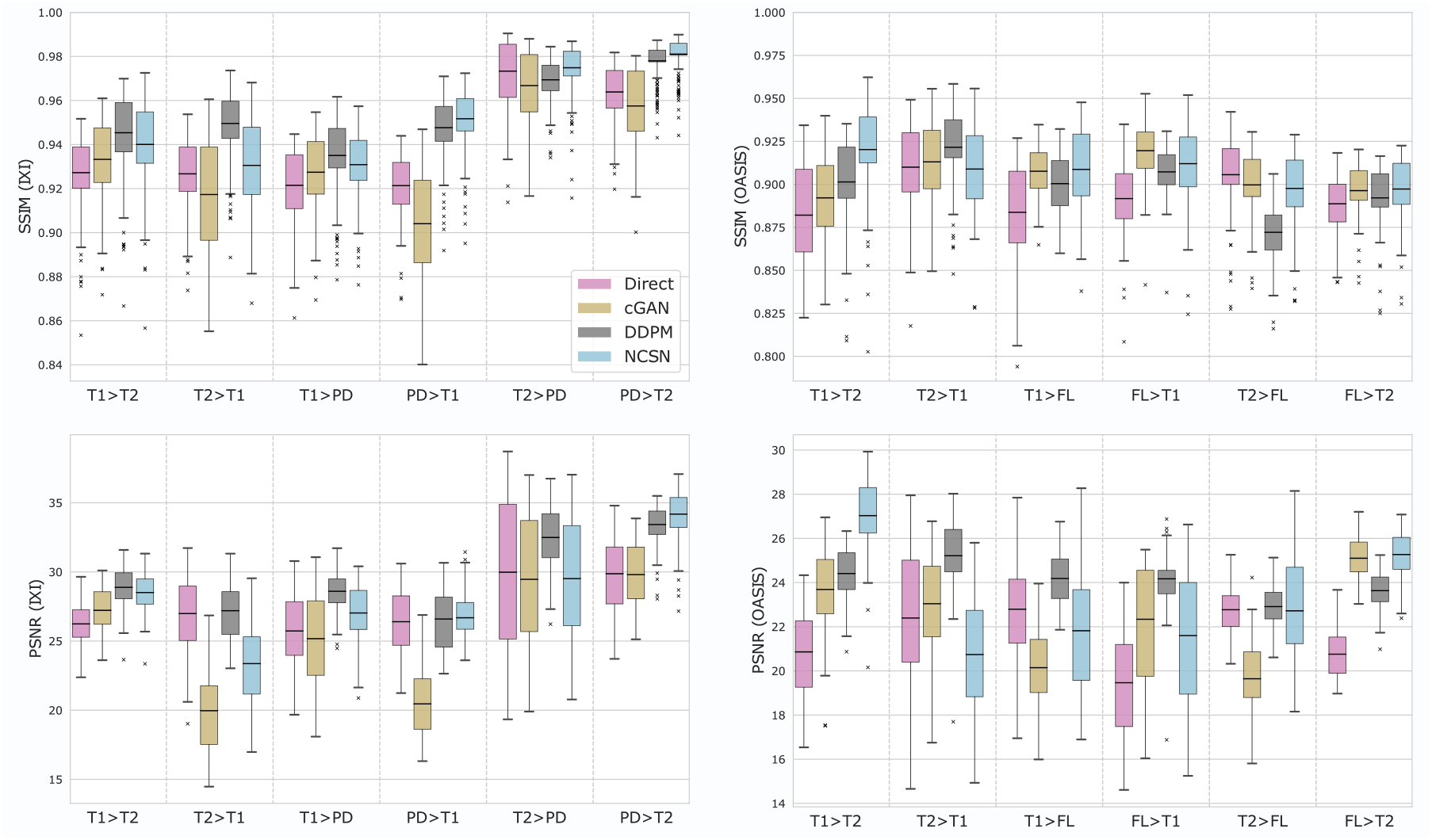
Boxplots SSIM and PSNR for IXI and OASIS datasets for Direct (pink), cGAN (brown), DDPM (gray), and NCSN (blue) models. Each group of four columns shows results for a given image transformation type.

In Table 1, we present the numerical values of the SSIM and PSNR metrics for the test subjects from the IXI and OASIS datasets. We dedicate four columns to the transformation models (Direct, cGAN, DDPM, and NCSN). Each row presents the numerical values (the average value and the standard deviation in parenthesis) for the specified transformation, where the metrics are calculated for all test subjects. For each row, we highlight the model with the highest SSIM. We use non-parametric Wilcoxon signed-rank test with *p <* 0.05 to assess the statistical significance of the calculated metrics. We observe that out of the twelve transformations, the NCSN model performs the best in six cases, the DDPM is the best in four cases, and the cGAN and Direct models are the best in one case each.

**Table 1:** Statistics of metrics (SSIM and PSNR) for test subjects from IXI and OASIS datasets for Direct, cGAN, DDPM, and NCSN models. Each row lists the metrics (mean and standard deviation in parenthesis) for a different transformation, with the best results highlighted in bold font.

| IXI | Direct<br>SSIM - PSNR | cGAN<br>SSIM - PSNR | DDPM<br>SSIM - PSNR | NCSN<br>SSIM - PSNR |
| --- | --- | --- | --- | --- |
| $T1 \rightarrow T2$ | 92.72(1.74) – 26.24(1.54) | 93.33(1.83) – 27.22(1.50) | **94.54(1.95) – 28.88(1.44)** | 94.01(2.00) – 28.51(1.30) |
| $T2 \rightarrow T1$ | 92.67(1.62) – 26.99(2.82) | 91.73(2.63) – 19.97(2.89) | **94.96(1.46) – 27.20(1.99)** | 93.05(2.17) – 23.37(2.91) |
| $T1 \rightarrow PD$ | 92.15(1.65) – 25.73(2.49) | 92.74(1.73) – 25.17(3.04) | **93.50(1.69) – 28.60(1.35)** | 93.09(1.55) – 27.03(2.02) |
| $PD \rightarrow T1$ | 92.13(1.41) – 26.40(2.36) | 90.41(2.27) – 20.46(2.50) | 94.77(1.39) – 26.59(2.05) | **95.17(1.36) – 26.68(1.47)** |
| $T2 \rightarrow PD$ | 97.33(1.59) – 29.98(5.34) | 96.68(1.63) – 29.47(4.47) | 96.94(0.96) – 32.49(2.26) | **97.49(1.17) – 29.52(4.11)** |
| $PD \rightarrow T2$ | 96.39(1.20) – 29.87(2.45) | 95.75(1.72) – 29.81(2.26) | 97.80(0.83) – 33.43(1.42) | **98.10(0.82) – 34.18(1.80)** |
| OASIS | Direct<br>SSIM - PSNR | cGAN<br>SSIM - PSNR | DDPM<br>SSIM - PSNR | NCSN<br>SSIM - PSNR |
| $T1 \rightarrow T2$ | 88.21(3.03) – 20.86(1.82) | 89.22(2.58) – 23.69(2.05) | 90.14(2.89) – 24.41(1.21) | **92.02(3.05) – 27.03(1.80)** |
| $T2 \rightarrow T1$ | 91.00(2.86) – 22.39(3.03) | 91.31(2.55) – 23.04(2.22) | **92.16(2.39) – 25.22(1.71)** | 90.89(2.81) – 20.74(2.54) |
| $T1 \rightarrow FL$ | 88.38(3.31) – 22.79(2.60) | 90.76(1.50) – 20.15(1.80) | 90.04(1.80) – 24.19(1.19) | **90.87(2.50) – 21.82(3.13)** |
| $FL \rightarrow T1$ | 89.17(2.51) – 19.47(2.39) | **91.96(1.96) – 22.34(2.61)** | 90.73(1.57) – 24.17(1.44) | 91.20(2.61) – 21.60(2.96) |
| $T2 \rightarrow FL$ | **90.56(2.70) – 22.77(1.12)** | 89.97(2.09) – 19.64(1.78) | 87.21(1.77) – 22.91(0.95) | 89.76(2.45) – 22.72(2.38) |
| $FL \rightarrow T2$ | 88.87(1.68) – 20.76(1.09) | 89.64(1.64) – 25.10(0.94) | 89.21(2.02) – 23.63(0.88) | **89.73(2.02) – 25.27(1.11)** |

When inspecting results from the two datasets, we observe that images from the IXI dataset are of better quality with less heterogeneity. On the other hand, images from the OASIS dataset are closer to clinical data but, in some cases, suffer from low contrast and artifacts. This is especially noticeable in Flair images from the OASIS dataset. This heterogeneity has resulted in lower metrics for images from the OASIS dataset when compared to their counterparts from the IXI dataset.

We also make several observations based on the visual inspection of the transformed images. For example, blurriness in the transformed images is more common for the Direct model. This likely stems from the application of the MSE (*L*_2_) error loss function, which pushes the pixel intensities of the outputs toward the target values but saturates when differences are minute. Notably, the Direct model outperforms the other models only in OASIS, T2 to Flair transformation, when the target images are inherently blurry, and the image types are close to each other. We also experimented with the mean absolute error (MAE or *L*_1_) loss function (not reported here for brevity) and found that it did not improve the overall performance.

When comparing the two diffusion models (which are also the two best-performing models), the DDPM output appears to be visually sharper. This feature is useful in applications where the high-frequency features of the output image, such as the texture, are important. However, as we mentioned before, the occasional presence of noisy voxels in the DDPM output can be misinterpreted as a signal (such as lesions in the Flair image). Therefore, depending on the downstream task, we recommend special attention to handling noise in the output. On the other hand, we did not observe any noise in the NCSN outputs that required extra post-processing.

In conclusion, among all models, the two diffusion models display better performance in MR image transformation tasks. Further, among the two diffusion models, the NCSN model is better both in terms of quantitative metrics and visual inspection of the results.

### 3.2 Role of probabilistic transformation

As discussed in 2.3, a critical aspect of the generative models is their ability to generate an ensemble of outputs for any input image. Here, we investigate the utility of generating this ensemble versus just a single output image. Notably, we demonstrate how the statistics of this ensemble respond to out-of-distribution (OOD) images that are used as input. We present these results in Figure 9, where the first row corresponds to a normal image sample, denoted as original, and the remaining rows contain OOD image inputs that are derived from this original image. The OOD images are synthesized by introducing common MRI artifacts to the original image.

**Figure 9:**
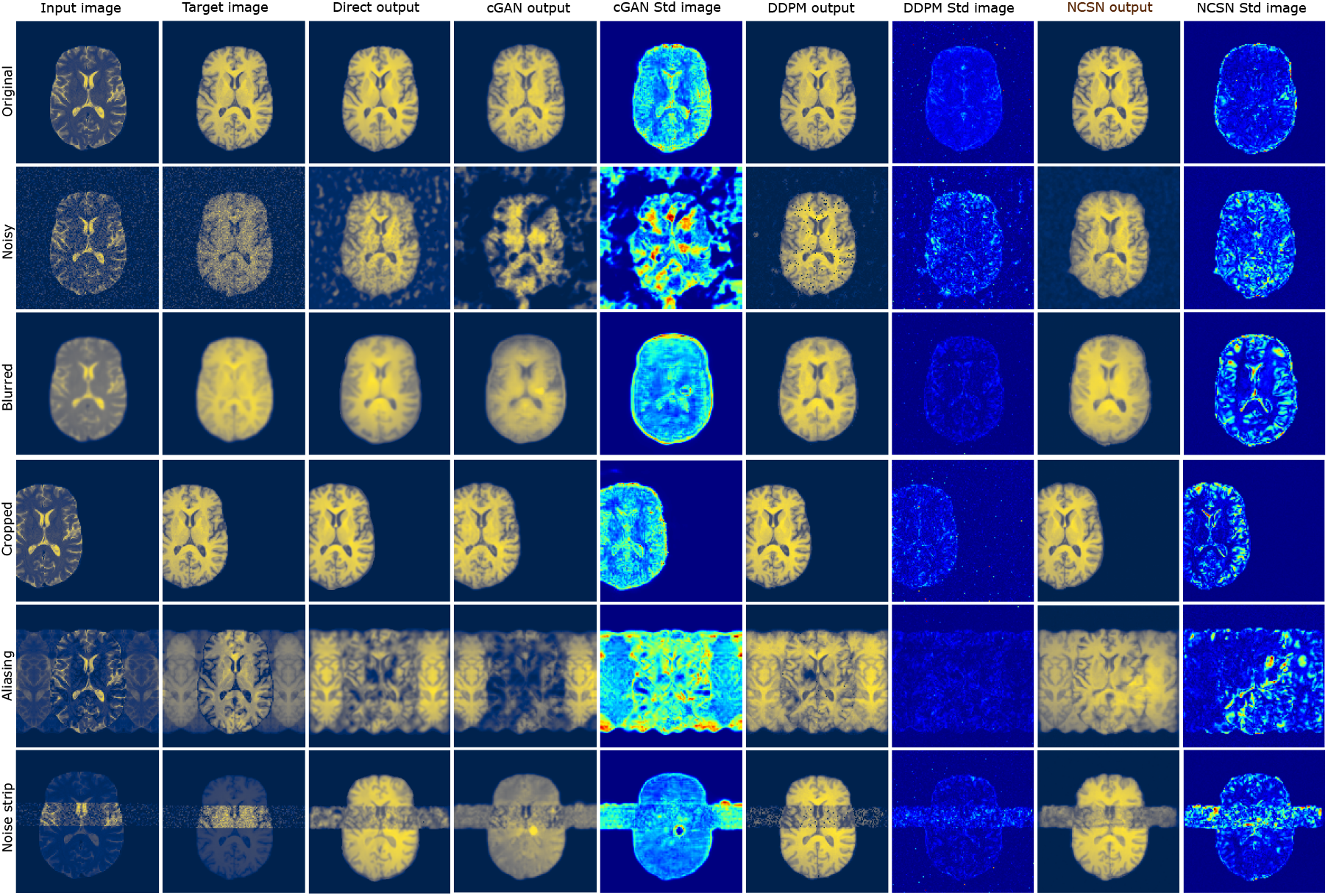
Response of models to typical and out-of-distribution (OOD) input images.Column 1: input image; column 2: target image; columns 3: output for the direct model; columns: 4-5: output and standard deviation for cGAN; columns: 6-7: output and standard deviation for DDPM; columns: 8-9: output and standard deviation for NCSN. Row 1: typical slice (Original); row 2: added noise artifact; row 3: blurring artifact; row 4: cropping artifact, row 5: aliasing artifact; row 6: noise-strip artifact.

In the second row, we represent an example of the original image altered by adding random Gaussian noise. We observe that the Direct and cGAN models fail to generate the brain boundaries and produce cloud-shaped outputs outside the brain in their prediction. Conversely, the geometry is less affected in the output of the diffusion models, especially the DDPM model. We also note that the DDPM model generates out-of-range spots that were discussed in Section 2.9. In the third row, we present the original image distorted by a blurring filter. Although the input is blurry, some models are successful in reconstructing sharper output images, which can be seen especially in the DDPM output. The fourth row demonstrates a cropped image. In this case, all models are able to handle this artifact effectively. The fifth row considers an image with an aliasing artifact, where we observe that all four models fail to preserve the geometry of the brain tissues. In the last row, the artifact is in the form of a noise strip, and once again, all four models are challenged by it. In summary, all four models are able to work with the cropped artifact, the diffusion models are better when working with noisy and blurred artifacts, and all four models fail when working with aliasing and noise-strip artifacts.

An advantage of the probabilistic models is that they allow the evaluation of the variance in the output which can be considered as a measure of uncertainty in the output. It is worth considering whether this measure of uncertainty is somehow correlated with the inputs that the model has not seen before - the OOD inputs. From Figure 9, that this is indeed the case, and that out of the three generative models, the NCSN model is most consistent in that it reports larger uncertainty when the model output is incorrect.

We now consider whether the standard deviation images in Figure 9 can be used to identify OOD input images as part of a quality control (QC) process. This is especially applicable in a clinical context where detecting corrupted data is critical in ensuring the reliability of the models before using their output in subsequent tasks. To that end, we first define a measure denoted as 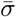 as the average of all voxel values in the three-dimensional standard deviation image of a subject. Next, we select five different artifact types (noisy, blurred, cropped, aliasing, and noise strip) and apply them to fifteen subjects from the test dataset, yielding a total of 75 corrupted 3D images. We use these, in addition to images from 144 normal subjects, as input to all three generative models. Thereafter, we plot the values for 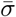 for each subject and model in the swarm plots in Figure 10. In each plot, the group of points to the right belong to normal images, while those on the left correspond to images with artifacts. Furthermore, we depict 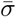 values for different artifact types with distinct colors.

**Figure 10:**
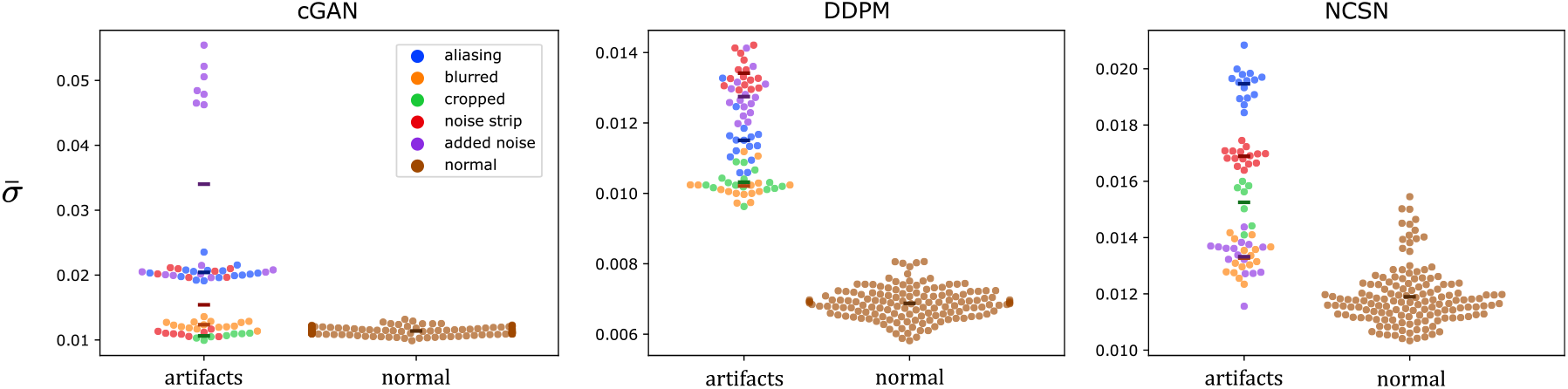
Distribution of 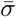 for cGAN, DDPM, and NCSN models. Distributions are divided into normal (right) and atypical input images (left), with artifact types indicated using different colors.

These plots can be used to answer two questions. The first is whether the value of 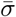 generated by each model can be used to distinguish OOD inputs from their normal counterparts. From the plots, we observe that while all three models display this characteristic to some extent, the DDPM is most effective. For this model, there is no overlap in the values of 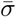 between normal and OOD images. Therefore, beyond the threshold of 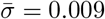 we can conclude that the input image is OOD. On the other hand, for the cGAN and NCSN models, we observe that there is some overlap in 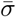 between normal and OOD images. The use of 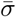 as a classifier to distinguish between normal and OOD images reveals that area under the receiver operator curve (AUROC) for the cGAN, NCSN, and DDPM models is 0.800, 0.946, and 1.00, respectively.

The second question is whether the value 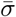 can be used to infer the type of OOD artifact. This would be true if different OOD types would cluster around specific values of 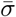. In this respect, we observe that NCSN model is most effective. For this model, the aliasing, noise-strip, cropped and blurred and noisy images tend to cluster into four distinct clusters.

## 4 Conclusions and Future Work

In this paper we have investigated the effectiveness of a class of deep learning-based methods called conditional generative models in transforming MR images. To that end, we implemented and evaluated the conditional version of three major generative models, namely, Conditional Generative Adversarial Network (cGAN), Denoising Diffusion Probabilistic Model (DDPM), and Noise Conditioned Score Network (NCSN). We also included a U-Net based deterministic model in our comparisons. We trained these models on the publicly available IXI and OASIS datasets and then applied the trained models to transform T1, T2, FLAIR, and PD images. We quantified this performance by computing image similarity metrics like SSIM and PSNR for the transformed and target image pairs. Through our comparisons, we concluded that the NCSN model demonstrated the best performance both in terms of quantitative metrics and based on visual inspection of the results.

We also investigated the utility of the generative property of the models, i.e., their ability to produce an ensemble of outputs for every input image. In particular, in addition to the standard MR images, we created an out-of-distribution (OOD) set of images by adding typical artifacts like blurring, aliasing, and Gaussian noise. We then used these images as input to the generative models, and for each input image generated an ensemble of sampled transformed images. Using these samples, we computed the average of voxel-wise standard deviation values for the regular and OOD datasets and determined whether this metric could be used to identify input images that were OOD. We discovered that the DDPM model was the most effective in this regard where a given threshold for this metric perfectly separated all the OOD input images from regular images. We believe that this ability will be particularly useful in the automated discovery of artifacts during the clinical application of these methods.

We also note that while the diffusion models (NCSN and DDPM) tended to perform better than the cGAN model, they are generally more computationally expensive. The DDPM and NCSN models require approximately 15 and 13 seconds, respectively, to generate a single slice using a NVIDIA Ampere Tesla A40 GPU with 48 GB memory. On the other hand, the cGAN and the U-Net models needed less than 0.1 seconds for the same task. However, we note that this drawback can be overcome using a multi-threaded architecture, where multiple slices can be generated in parallel.

In this work, we have focused on implementing and quantifying the performance of the proposed models in terms of image similarity metrics. A potential follow-up study could explore the use of transformed images in downstream tasks. For instance, most structural analysis software, like FreeSurfer, primarily rely on T1 MR scans. In the follow-up study, one could consider the use of native T1 images, and T1 images obtained from transforming T2 images, in FreeSurfer and compare the resulting outputs from the two sets. The outputs would include quantities of clinical interest, such as the volume of brain tissue and/or lesions. An analysis of this type would also provide the groundwork for determining whether MR images of a given type, such as T2 images, can effectively replace T1 images in these applications.

## Appendix

### A- Network Architectures and Hyperparameters

We aimed to keep the hyper-parameters consistent across the models whenever possible while adhering to the original implementation of each work. Accordingly, we used the same training and testing data for all models with an image resolution of 128 × 128 and a batch size of 16. The learning rate is set to 1*e* −4 with Adam as the optimizer and amsgrad [47] option enabled in all training experiments. As mentioned earlier, the neural network architecture details for the cGAN critic and generator with conditional instance normalization (CIN) [48] can be found in [32] and [27]. We replaced the ResBlock of the generator U-Net with a Dense block discussed in [49]. Also, the details of the U-Net for the DDPM can be found in [36] and [37], and for the NCSN models in [34] and [35]. We note that we adopted the U-Net architecture used in the DDPM model for the Direct model by eliminating the time embedding layers. The number of channels for the first level in the U-Nets is chosen to be 64 for all models, resulting in 20 ± 2 million trainable parameters (weights) for all models, except for the cGAN critic which has 5.8 million weights. For both diffusion models (DDPM and NCSN), we used 1000 diffusion steps. As previously mentioned, *σ*_*t*_ are learned during the training by the model in the DDPM method. For NCSN, the authors propose a technique to determine the initial noise scale.

They suggest setting the largest noise scale to be as large as the maximum pairwise Euclidean distance of training data points. We note that this value differs depending on the dataset and should be calculated based on the output image type dataset (as the noising process is applied to the output image). Also, using the maximum Euclidean norm of all data points can be a resonable estimate for the maximum noise scale in MR images. For all models, we used 0.01 as the minimum noise scale.

## 5 Acknowledgments

A.A.O. would like to acknowledge the support of ARO Grant AWD-00009144.

C.J.A. receives grant support from the National MS Society, the National Institutes of Health, and the Keck School of Medicine. She has received consulting honoraria for participation in advisory boards, data safety monitoring committee, and/or MRI Adjudication Committee for Horizon Therapeutics, TG Therapeutics, EMD Serono, Genentech, and Zenas Biopharma; and for participation in educational events for Sanofi Genzyme, MJH Life Sciences, Efficient LLC, and Spire Learning

The authors acknowledge the Center for Advanced Research Computing (CARC) at the University of Southern California for providing computing resources that have contributed to the research results reported within this publication. URL: https://carc.usc.edu.

